# Viral Taxonomy Derived From Evolutionary Genome Relationships

**DOI:** 10.1101/322511

**Authors:** Tyler J. Dougan, Stephen R. Quake

## Abstract

We describe a new genome alignment-based model for classification of viruses based on evolutionary genetic relationships. This approach uses information theory and a physical model to determine the information shared by the genes in two genomes. Pairwise comparisons of genes from the viruses are created from alignments using NCBI BLAST, and their match scores are combined to produce a metric between genomes, which is in turn used to determine a global classification using the 5,817 viruses on RefSeq. In cases where there is no measurable alignment between any genes, the method falls back to a coarser measure of genome relationship: the mutual information of k-mer frequency. This results in a principled model which depends only on the genome sequence, which captures many interesting relationships between viral families, and which creates clusters which correlate well with both the Baltimore and ICTV classifications. The incremental computational cost of classifying a novel virus is low and therefore newly discovered viruses can be quickly identified and classified.

## 1. Introduction

Collectively, viruses display an unstructured diversity which hinders the imposition of any systematic classification. Because viruses evolve quickly and lack an analogue to the bacterial 16S sequence, there is no consensus surrounding their taxonomy. The Baltimore classification groups viruses into seven categories based on the biochemistry of their replication strategies, nucleotide character, but as the basis for a phylogeny, it conflicts with the observation that some viruses with similar functions and structural proteins have different types of genomes.[1] The International Committee on the Taxonomy of Viruses (ICTV) uses a hierarchical taxonomy inspired by the modern tree of life. However, because this classification has been built up over time in order to accommodate new discoveries, it enjoys neither the tree of life’s coherence nor its authority.[2] When a new virus is discovered, subjective analysis is required in order to incorporate it into the taxonomy. Other classifications have been proposed based on structure, host species, or genome length.[16]

The viral genome captures a record of fingerprints of the evolutionary history of the virus and should in principle provide the basis for calculating relationships between any set of viruses. Hendrix et al. have shown that the genome encodes history including recombination events and virus-species coevolution.[17] The use of full genomes in viral classification, now possible because of modern sequencing techniques, offers the promise of weighing all salient features of a given virus more objectively than the ICTV’s consensus system. As of this writing, there are 5,817 full viral genomes on RefSeq[3] and in recent years several researchers have proposed alignment-free methods due to the computational complexity of sequence alignment and the rate at which new viruses are sequenced. Typically a vector in a Euclidean “genome space” is calculated from various properties of the genome, and distances between such vectors are used for classification. These include Yu et al. (2013) and Hoang et al. (2015).[4][5] Although they provide an injective mapping onto the chosen genome space, it is not clear that the distance between two points in genome space is a meaningful metric of genome similarity.

Alignment-based classifications, such as the one which will be described below, stand on stronger interpretive footing: the “distance” between two genomes can be defined directly from a similarity score returned by an alignment. Gene alignment is most appropriate for sequences that are very similar, and as we will see, a global classification can be built up by considering only these relationships. To find these relationships, we use the sequence alignment tool BLAST; its efficiency not only makes the alignment of 5,817 genomes tractable, but also reduces the laboriousness of re-computing the taxonomy after a significant number of new viruses have been sequenced. Rohwer et al. (2002) propose a phylogeny for phage using BLAST hits; although successful within this more limited scope, their heuristic distance metric makes binary distinctions, reducing the amount of information used.[15] Here, we present the results of a statistically motivated alignment-based classification which considers genomes as collections of individual genes; we find that it correlates well with the ICTV, host kingdom, and Baltimore classifications, and also provides additional insights beyond these.

## 2. Methods

Intuitively, we want to impose a distance metric on the space of viral genomes. From this distance metric, we should be able to extract clustering information, and build a taxonomy. We have sought to find a metric which captures the shared information between genes in a pair of genomes. This approach was inspired by the notion of mutual information from information theory, but uses a more simple measure of shared information based on current flow in parallel resistor networks. In this case information is meant to play the role of current (or, more precisely, information lost in evolution is meant to play the role of resistance) and the more genes that have relationships and the stronger the relationships then the stronger the overall relationship between the genomes is judged to be.

### 2.1 Gene Alignment Distance

Functionally, a genome is a collection of genes. This suggests that we begin calculating a distance between genomes by calculating the distances between their component genes. Suppose we have some way to quantify the dissimilarities between the genes in two viruses, and wish to calculate an overall distance between the viruses as a whole. The overall genome distance between virus A and virus B should satisfy certain properties. If all of the genes in virus A are completely different from all of the genes in virus B, their overall distance should be very large. If virus A and virus B are each composed of a single gene, their overall distance should be equal to the dissimilarity between these two genes. Each additional gene on virus A which is similar to a gene on virus B should only decrease, never increase, the total distance between the viruses.

The above properties are satisfied by the physics of resistors in parallel, suggesting that we calculate the total inverse distance by adding the inverses of the constituent dissimilarities. This analogy is shown in Figure 1: if we imagine a resistive wire connecting each of the genes for which a match is found, with resistance equal to the dissimilarity, then the total distance between the genomes is equal to the equivalent resistance between the two. Mathematically, the total distance *D*_eq_ between genomes *A* and *B* with genes *A*_*i*_ and *B*_*j*_ is 
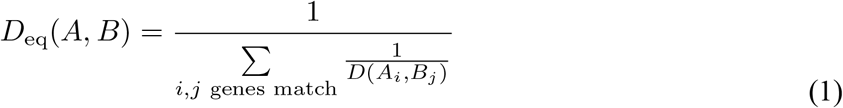
So given a suitable method for calculating the dissimilarity *D*(*A*_*i*_, *B*_*j*_) between two genes, we can calculate a meaningful distance between two genomes. All that remains is the calculation of dissimilarity between two genes, which requires a measurement of mutual information – a topic which has been well explored as part of the field of information theory.[12] Mutual information, a rigorous measure for the information shared between two variables, is defined as 
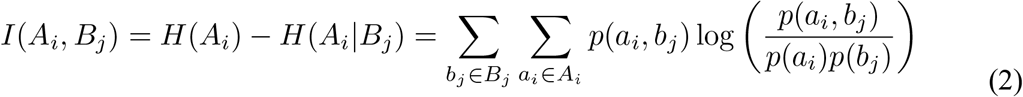
for two random variables (here, defined by genes) *A*_*i*_ and *B*_*j*_ which take values *a*_*i*_ and *b*_*j* (*i* and *j* index the genes, and play no part in the equation; they are retained here only for notational consistency). *I*(*A*_*i*_, *B*_*j*_) is the mutual information between genes *A*_*i*_ and *B*_*j*_, and *H*(*A*_*i*_) and *H*(*A*_*i*_ | *B*_*j*_) are the total and conditional Shannon entropies of genes *A*_*i*_ and *A*_*i*_ given *B*_*j*_, given by 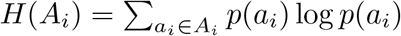 in analogy with_thermodynamic entropy.[12] Then 
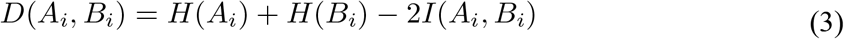
induces a distance metric, known as the variation of information. We will use this metric for our dissimilarity scores between two genes.

**Figure 1.**
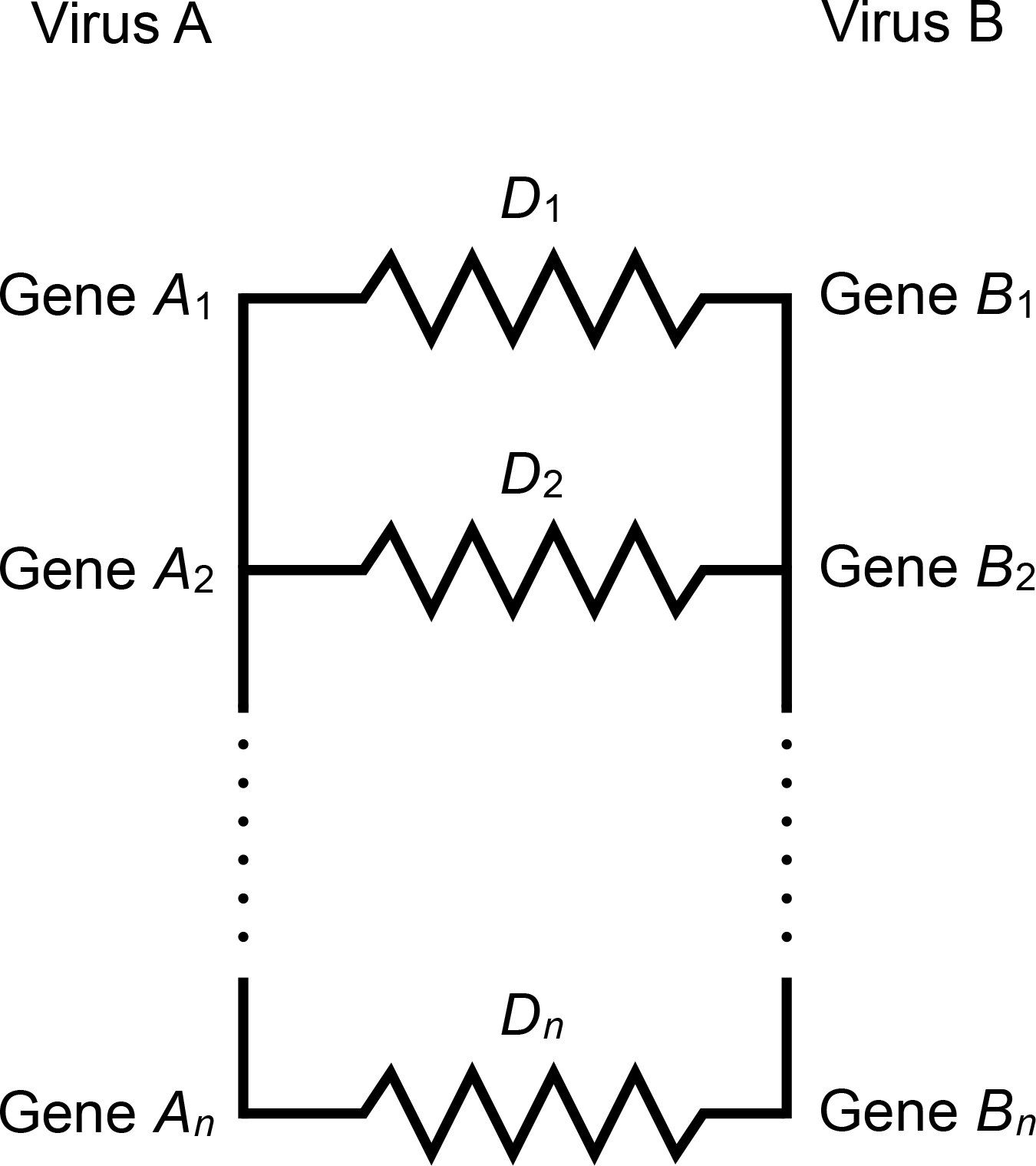
Graphical representation of “distance” calculation for two viruses. Each virus is made up of genes, some of which may match to the genes on another virus (Viruses A and B, respectively). We imagine the variation of information between two genes D_i_ as the resistance of a resistor connecting them. Then the total “distance” between the collections of genes in Viruses A and B is the equivalent resistance between the two sides. Only the genes A_i_ and B_i_ which match are shown and indexed; the total number of options for choosing one gene from each virus is quite large, and most such choices do not yield a match. These pairs are analogous to open circuits, with infinite resistance, which do not affect the equivalent resistance.

Intuitively, the mutual information between two genes should have several features. Two genes which are identical should have a mutual information proportional to their length. Two genes which share nothing in common should have zero mutual information. Each additional bit of mutual information between two genes should represent one additional binary choice which can be correctly determined about one gene, knowing the other. Described another way, each additional bit should represent a doubling of the possible set of random genes from which one gene could be discerned, given knowledge of the other gene.

To calculate the mutual information between genes, we perform sequence alignment using BLAST. We treat each gene as a separate sequence, which results in a database of 300,000 sequences. For each gene, we attempt an alignment of the translated protein sequences against every other sequence using BLAST (TBLASTX). BLAST searches for short match regions between two sequences, and then lengthens these matches to find regions of similarity. Because of a series of simplifications, BLAST is one of the most computationally efficient alignment algorithms available. The BLAST search returns various statistics on the quality of each match; of these, we will concern ourselves with the bit score. For a fixed database, the bit score is a logarithmic function of the e-value, and is linearly related to both the percent match and the alignment length. Moreover, the bit score has a convenient interpretation: it is the size, in bits, of a database one would need to search through to find an equally good match by chance.

It then makes intuitive sense that the bit score should be proportional to mutual information, and indeed this has been shown by Mazandu et al. (2011).[6] For a fixed database, this proportionality constant varies little between searches; see Korf et al. (2003) for details.[7] Our analysis is unchanged by an overall scaling factor, but this proportionality constant will become important later on. The entropies are determined by the mutual information between a gene and itself. If the mutual information is the amount of information shared between two genes, then the variation of information is the amount of information contained in each virus which is not shared by the other.

Now, given two genes which are found by BLAST to match, we can calculate their dissimilarity using the variation of information; given several such matches between two viruses, we can calculate the distance between the viruses. In the resistor model, gene pairs for which no match is found may be thought of as wires with infinite resistance, or open circuits in parallel.

This formula was used to calculate a distance matrix for all of the viruses considered. Defined distances were found for 30% of the pairs of viruses, and the other 70% of pairs had no matches, because all of their genes were divergent from each other. This was to be expected; alignment fails on highly divergent sequences. Nonetheless, when considering the viruses individually, 70% of the viruses had a match with at least one other virus. Motivated by the need to classify the other 30% of viruses, for which no gene matched any other gene in a virus, we turn to a different genome property from which to calculate a variation of information: k-mer frequency. Ultimately, these two metrics will be combined in order to produce a single comparison between all viruses.

### 2.2. K-Mer Distance

It is also possible to calculate the variation of information based on k-mer frequency in genomes, which returns a finite value for all pairs of genomes. K-mer frequency analysis has been a widely applied and useful approach in microbial genome sequencing and in metagenomic analysis.[13-14] This is a cruder metric than the BLAST based approach described above, but it enables one to compute mutual information in cases where there are no gene alignments between genomes.

We compare the prevalence of 256 possible k-mers of four letters in each genome. Consider each possible k-mer as an outcome in the probability theory sense, and build a sigma-algebra out of the power set of k-mers, so that each event is a unique set of k-mers between 0 and 256 in size. Define a probability measure on this sigma-algebra which simply gives the fraction of possible 4-mers included in the event, so for example, P({AAAA, ATAT, ATCG}) = 3/256 because this event encompasses three k-mers. This gives a probability space. Finally, note that in a given genome, each k-mer may be more or less prevalent than it is on average in all genetic material in the data set. For a given virus, define a random variable *A* whose input is a given k-mer and whose output is whether it is over-represented or under-represented in that genome. The random variable *A* can take two possible values *a*: under-representation and over-representation. The probability of over-representation is the fraction of the 256 k-mers which are more prevalent in the given genome than they are in the dataset at large.

This probability space allows for direct calculation of variation of information between two genomes. The mutual information is given by 
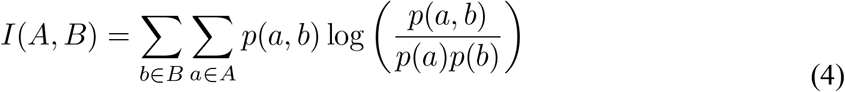
If, for example, *a* = *b* = k-mer over-representation, then p(x,y) is the fraction of k-mers which are over-expressed in both viruses. If two viruses have similar distributions of k-mers, then the joint probabilities of expression in both genomes will be very high; if they have dissimilar distributions, then representation in one virus will be largely independent of representation in the other, leading to no mutual information. Variation of information can then be calculated as above, providing a finite distance between every two genomes in the data set. Using 4-mers gives a distribution of variation of information heavily skewed away from 0.

To determine how this distance should be combined with the gene alignment distance above, we note a few general facts. Genes carry the most salient and most conserved information in genomes, so gene alignment, when present, should be considered a better determinant of similarity than k-mer frequency. K-mer frequency is most relevant when two genomes have no BLAST matches, but for continuity, should contribute to all distances. Because k-mer frequency dissimilarity can be thought of as almost independent of gene alignment dissimilarity---large portions of each genome do not code for proteins, and these can be the ones thought of as setting the k-mer frequencies---the simplest solution is to add the inverse of the k-mer dissimilarity to the inverse of the gene alignment dissimilarity, effectively treating it as another resistor in parallel.

There remains a problem of magnitudes; the gene alignment distances which are finite range from almost zero to almost 30,000 bits, while the values for the k-mer variation of information range between 0 and 2 bits. In order to prevent the k-mer distance from eclipsing the gene alignment distance in all but a few cases (as a 30 kΩ resistor would become irrelevant in parallel with a 2Ω resistor), the two distances must be scaled relative to one another. In order to set similar scales between the two distances, we rescale the k-mer variation of information so that its maximum is equal to the maximum gene alignment distance; the very strong left skew in the k-mer distance ensures that when the gene alignment distance is finite, gene alignment dominates in all but 5 cases out of 10,000,000; in these 5, the two distances are comparable.

Although we have argued that the two metrics attempt to measure independent genome features, a sanity check is provided by their consistency. The distances are not directly correlated---most k-mer distances are near 2, and most gene alignment distances are near 0---but the metrics are not independent. In particular, for a given gene alignment distance, there is a lower bound on the k-mer distance which increases monotonically. Clustering on only one of the two distances reproduces many of the same global features as clustering on the other, but they provide resolution in complementary areas; using only one but not the other is substantially less effective, as shown in Supplementary Figures 1 and 2.

### 2.3 Building a Phylogeny

The above methods produce a distance matrix, but further statistical procedures are needed before it can be used for a phylogeny. BLAST’s efficiency relies upon several heuristics, and in some cases, these heuristics lead to unphysical situations, such as negative dissimilarities when two different sequences match better with each other than they do with themselves. These were dealt with as simply as possible, e.g. by raising negative values to small positive values, in order to avoid dividing by zero. A further issue was that because of BLAST’s heuristics and the derived nature of the distance metric used, the distance matrix does not satisfy the triangle inequality, and is overall very noisy. Because any further clustering relies on the genome space being Hausdorff, we used classical multidimensional scaling to embed the genome space in 50-dimensional Euclidean space.

In order to further reduce noise while preserving as much structure as possible, we used nonlinear dimensionality reduction. This is because the higher-dimensional spaces from which it is derived are too noisy to extract the salient features of the space. The data showed evidence of nontrivial topologies in two dimensions, so all data analysis was performed on a three-dimensional embedding, which is not shown. (For example, note that in Figure 4, cluster 16 is disconnected, so although it is shown to be a single cluster by the clustering in three dimensions, it resembles a torus, so a two-dimensional image of it is inadequate.) Because our alignment-based method produces more significant results for viruses which are quite similar, we chose to use t-SNE dimensionality reduction, which preserves local structures more than global ones. A Barnes-Hut implementation of t-SNE was used, with perplexity 30 and ¸ = 0.5. The algorithm was run 20 times, and the result with the lowest Kullback-Leibler divergence was chosen. This produced clear clusters of various shapes and sizes. Because t-SNE attempts to preserve neighbor probabilities, it suggests a density-based clustering algorithm; for the analysis, we used OPTICS, a variant of DBSCAN which produces hierarchical clustering on the three-dimensional space. This produced results almost identical to those of nearest-neighbor clustering.

## 3. Discussion

Figure 2 gives a 2-dimensional t-SNE visualization of the 50-dimensional data, using initial values from a 2-dimensional PCA projection of the 3-dimensional space used for clustering. These are colored by the 78 clusters found in the 3-dimensional t-SNE space. The number of clusters was relatively stable at 78, i.e. a relatively wide range of tree cuts all produced 78 clusters. This is comparable to the 125 families determined by ICTV. Most clusters consist of a single Baltimore type, as shown in Figure 3. Similar results arise when the data are colored by host kingdom instead, as shown in Figure 4. In Figure 5, these clusters are shown as a dendrogram.

**Figure 2.**
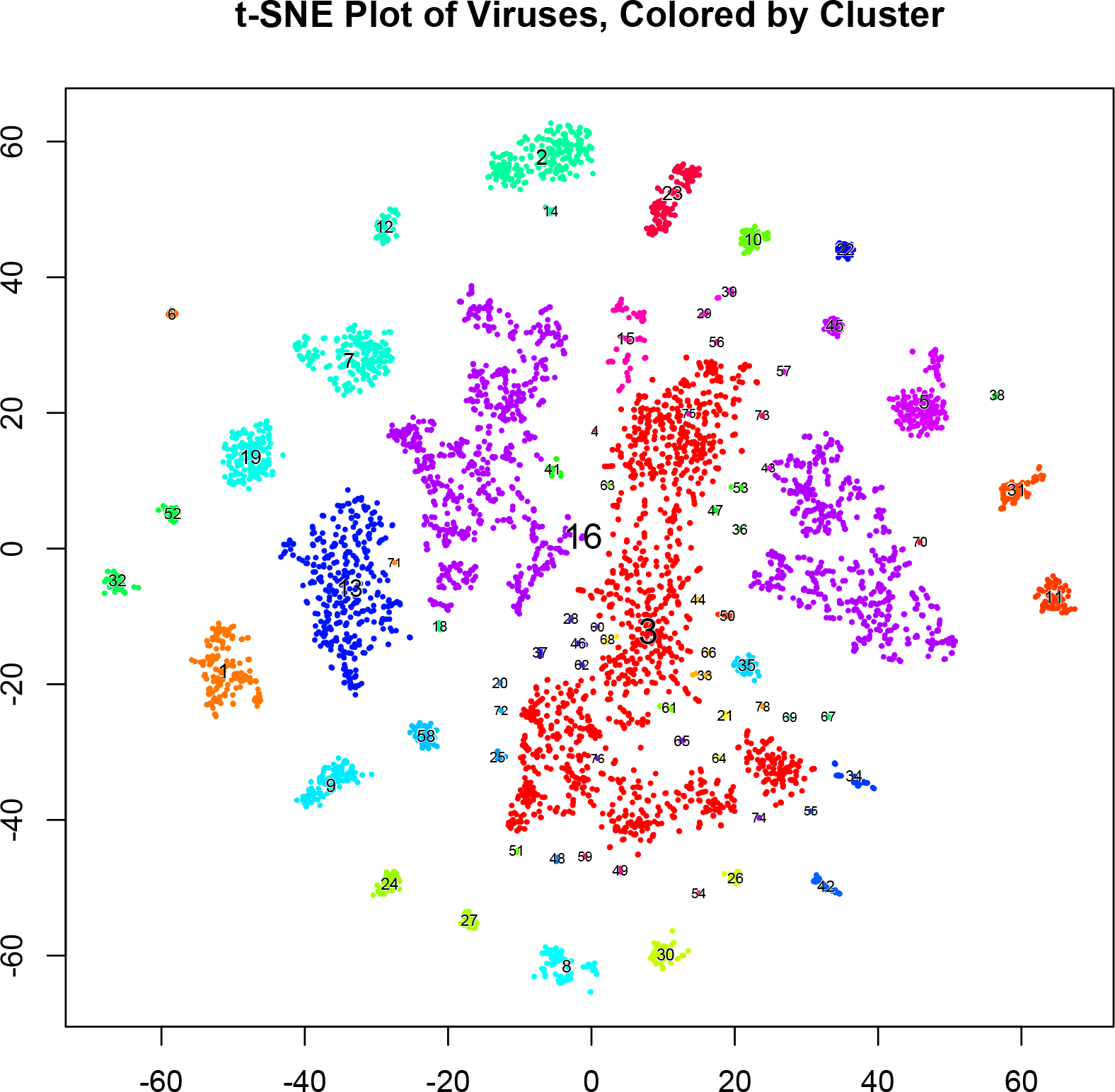
t-SNE plot of 5,817 viruses from RefSeq, grouped by mutual information of genes in common and k-mer frequency. Points are colored according to 78 clusters assigned by density-based clustering in the 3-dimensional t-SNE space. Each cluster is numbered, and cluster numbers correspond to those in Figure 5. (Note: the two large blue regions in the center are both part of cluster 16, even though they are not connected in the 2-dimensional space.) Most clusters correspond to one order or family in the ICTV classification.

**Figure 3.**
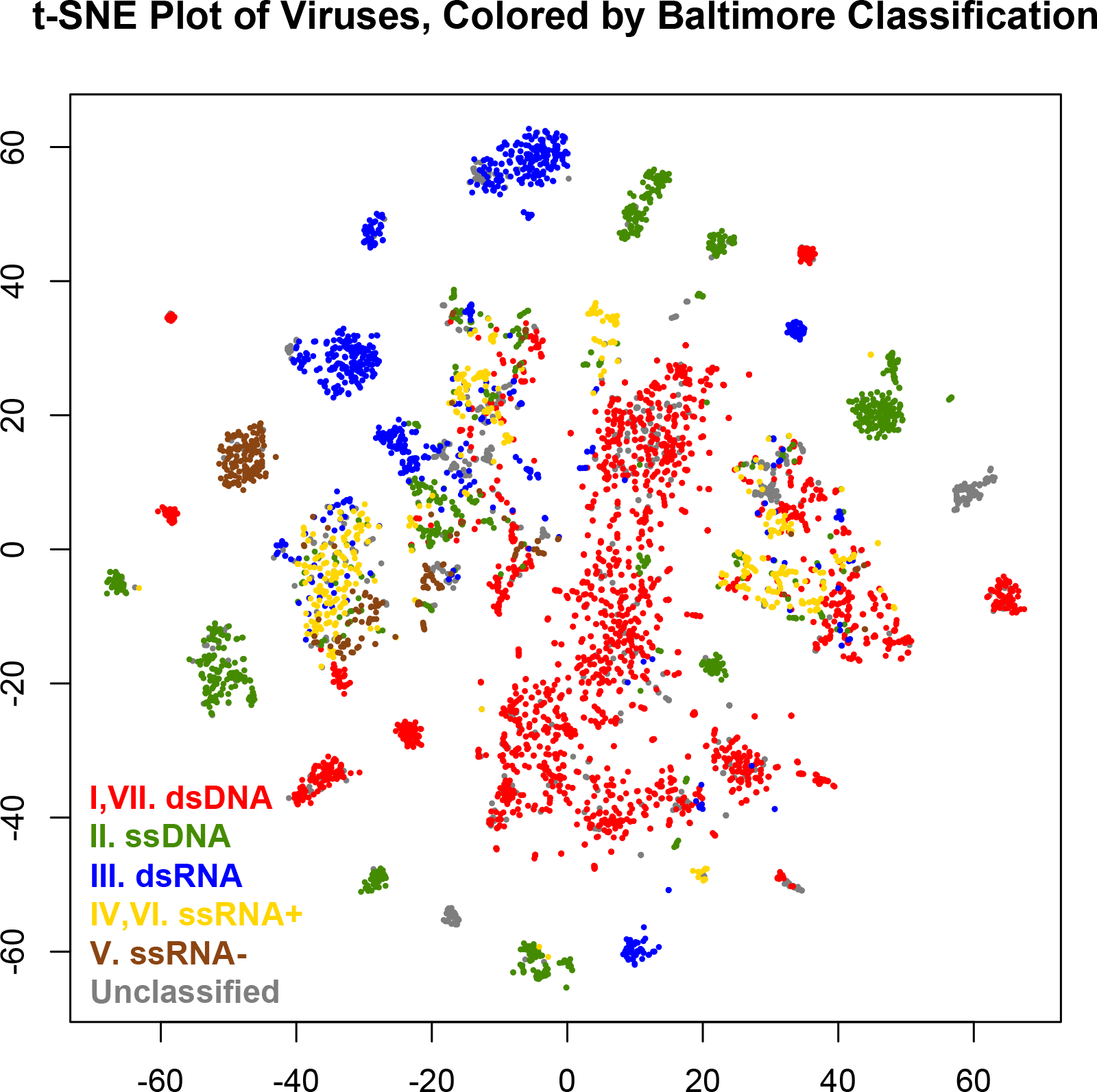
t-SNE plot of 5,817 viruses from RefSeq, grouped by mutual information of genes in common and k-mer frequency. Points are colored by Baltimore classification: red = dsDNA viruses (Baltimore class I & VII), green = ssDNA viruses (Baltimore class II), blue = dsRNA viruses (Baltimore class III), yellow = ssRNA viruses, positive sense (Baltimore class IV & VI), brown = ssRNA viruses, negative sense (Baltimore class V), and gray = unclassified.

**Figure 4.**
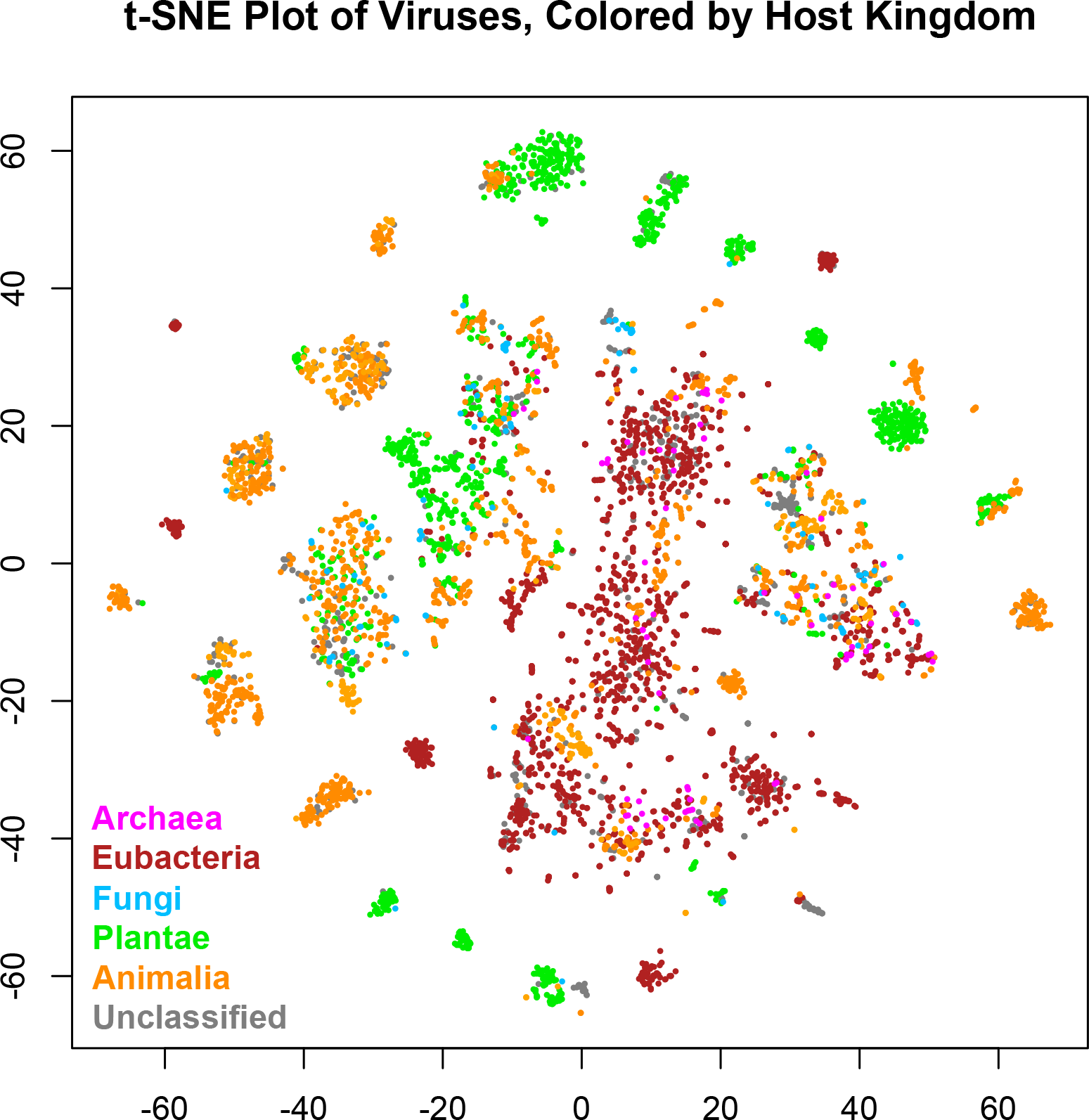
t-SNE plot of 5,817 viruses from RefSeq, grouped by mutual information of genes in common and k-mer frequency. Points are colored by host kingdom: magenta = Archaea, maroon = Eubacteria, sky blue = Fungi, lime green = Plantae, orange = Animalia, and gray = unclassified.

**Figure 5.**
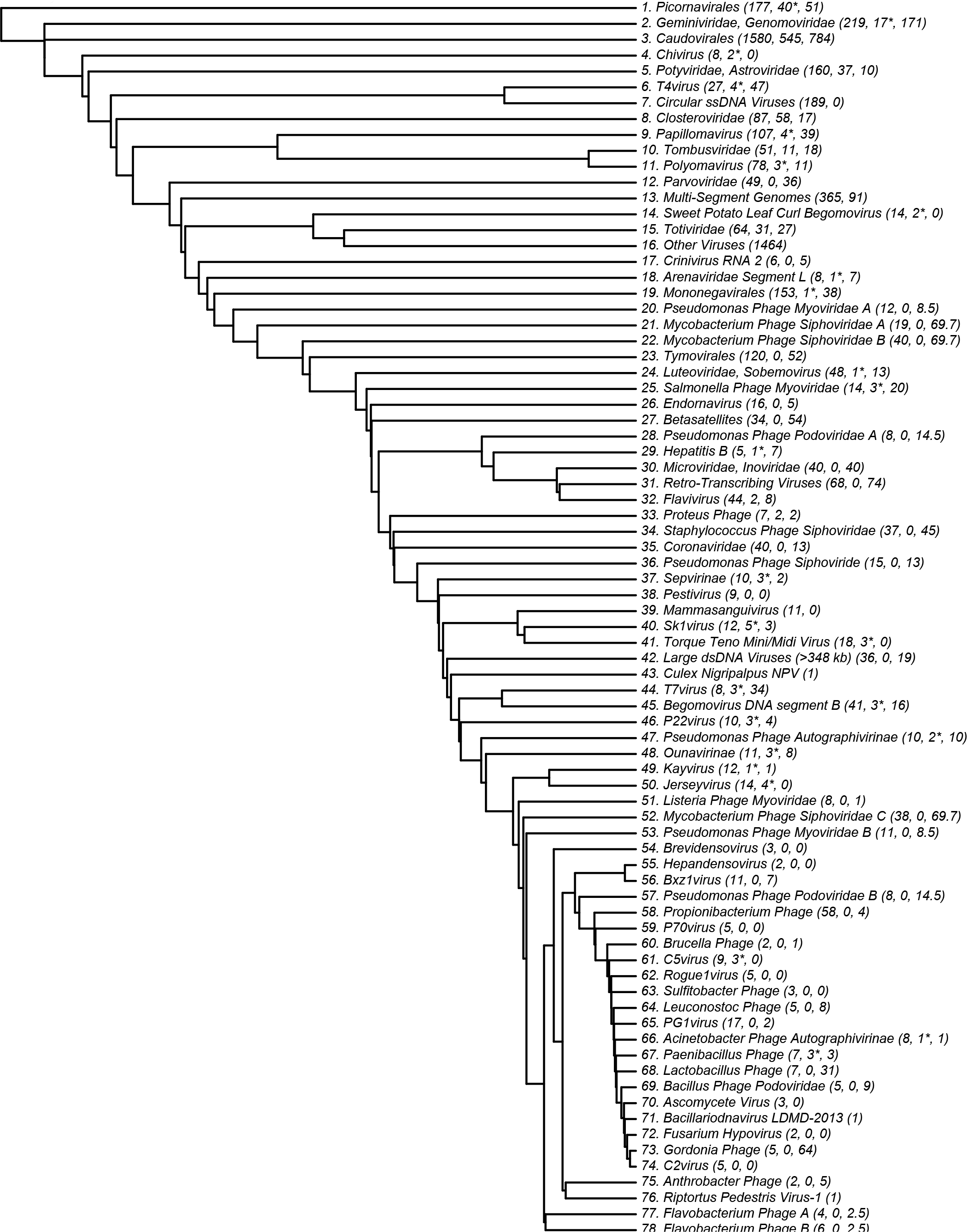
Dendrogram of 78 clusters of viruses, determined by density-based clustering on the 3-dimensional t-SNE space of viral genomes. Cluster numbers correspond to numbers in Figure 4. Three numbers are given in parentheses: the number of viruses in each cluster, the number of viruses which are not part of the family but nevertheless are clustered together with it, and the number of viruses in a family which were not included in its cluster. Asterisks after the second number indicate that most of the incorrect viruses are unclassified by ICTV or otherwise correspond weakly with it.

Almost all clusters could be named using an established ICTV order, family, or genus. Interestingly, cluster 39 was not: it contains a collection of viruses which infect through mammalian blood, such as the Abelson murine leukemia virus, the Hepatitis C viruses, and Pegiviruses. We have named this cluster “Mammasanguiviridae.”

While the Hepatitis C viruses were found in cluster 39, the Hepatitis B viruses displayed significantly broader genetic diversity. Cluster 29 contained all four Hepatitis B viruses which infect bats or monkeys, as well as the bluegill hepadnavirus, an unclassified hepadnavirus which might be considered a Hepatitis B variant with fish as hosts. The strains of Hepatitis B with other hosts were found in various other clusters, predominantly in subcluster 31 of cluster 16. This suggests that for some families of viruses, the specific host may play an important role in determining the virus’s genetics.

While most clusters were mostly or completely unified by a single ICTV class or other feature, cluster 16 (Other Viruses) was not. In three dimensions, cluster 16 forms a ring around cluster 3, suggesting that it may carry a more complicated structure that is not well-represented in the 2-dimensional plot. In order to determine the organization of cluster 16, it is necessary to explore further out on the derived phylogenetic tree. This is done in Supplementary Figure 3, which displays a phylogenetic tree of 98 sub-clusters of cluster 16. Distances between viruses in cluster 16 are determined almost completely by the k-mer variation of information, which yields less conclusive relationships. Most sub-clusters in cluster 16 consist primarily or wholly of one ICTV class, but few contain every element in that class. A few clusters displayed no clear unifying characteristics. The algorithm also tended to elide viroids, satellites, and individual segments of multi-segment genomes; the clusters which lacked unity had large populations of such viruses.

Many families of viruses were assigned to clusters consistent with ICTV nomenclature. Of the viruses not classed with cluster 16, 80% were consistently clustered; most of the disagreements (75%) occurred in clusters 3 and 14. As an example, 42 out of 50 of the flavivirus family were found in cluster 12. Of particular interest is the Zika virus and its relation to the Dengue viruses. Traditionally, Zika has been grouped with Spondweni, and believed to be less similar to the four strains of Dengue than they are to each other.[11] However, recent discussion has questioned whether Zika may be more related to some of the Dengue viruses, and whether the differences between the Dengue strains may be more significant.[10] Our analysis is consistent with the traditional approach, as shown in Figure 6. We find Zika to be most genetically similar to the West Nile virus, yellow fever, and Spondweni, than to any of the Dengue viruses. Nevertheless, our findings affirm the significant heterogeneity between the Dengue viruses; only Dengue 1 and 3 are particularly similar.

**Figure 6.**
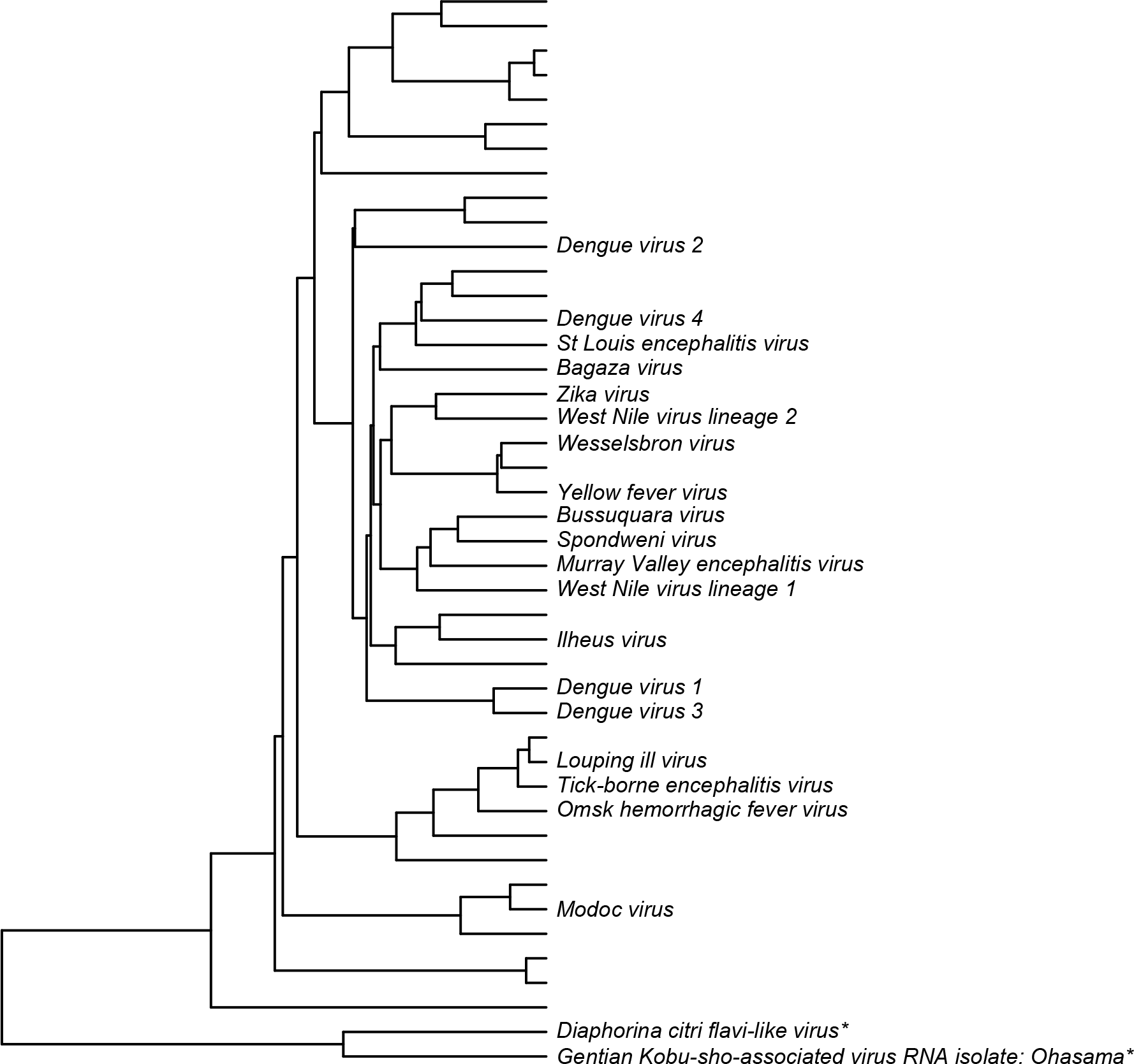
Dendrogram of cluster 32, which corresponds to the Flaviviruses. Of the 44 viruses in this cluster, 19 which are clinically significant in humans are labeled. This constructed phylogeny suggests that Zika is more genetically similar to the West Nile virus and yellow fever than to the Dengue viruses; the closest Dengue virus is Dengue 4. The two species marked with asterisks are not Flaviviruses; although they were clustered together with them, they are separated from the true Flaviviruses. Eight Flaviviruses make up a single sub-cluster of cluster 16 instead.

Because of their unique biochemistry, the genetic unity of the retroviruses is also worthy of note. Cluster 31 is made up wholly of retroviruses, suggesting that there are indeed some invariant patterns of expression associated with their transcription process. However, cluster 31 contains only half of the retroviruses in the study; the others are found in several different clusters, such as subcluster 35 of cluster 16, which contains four Polyomaviridae and three retroviruses, all with animals as natural hosts. Unlike most classifications, which rely upon one feature, our taxonomy incorporates predictors of size, biochemistry, host, and other factors, so the final classification can only be explained by a combination of these.

Although our model successfully produces a meaningful classification of viruses using only quantitative features of their genomes, it has certain limitations as well. BLAST is one of the only alignment tools fast enough to use for a global classification such as this, but its speed comes at the cost of many heuristics which diminish its accuracy. Our mutual information calculation, and especially our dimensionality reduction and clustering, are able to remove much of the noise in the raw BLAST data. Nevertheless, BLAST’s performance may limit the precision of our final classification and it would be of interest to explore whether more computationally intensive algorithms improve classification.

Our model is well-suited to partition the viruses into groups whose size approximates that of families. Because our model was developed for fidelity at great distances, it may be well complemented by other less heuristic alignment-based methods for finer scale analysis of very closely related genomes.

In conclusion, we have shown that a genetic classification using only quantitative features can provide a meaningful viral taxonomy. This taxonomy not only corroborates other classification systems, but also can be used as a check on them. Incremental viruses could be added without having to re-compute the whole structure, because they would not change it measurably, and even the entire analysis only takes a few days on standard computing resources. Future work could focus on improving the speed and accuracy of alignment tools. However, the search for a classification of viruses will be most aided by the sequencing of new viruses, in order to better understand the structure of the phylogenetic tree as a whole.

---

**Supplementary Figure 1.**
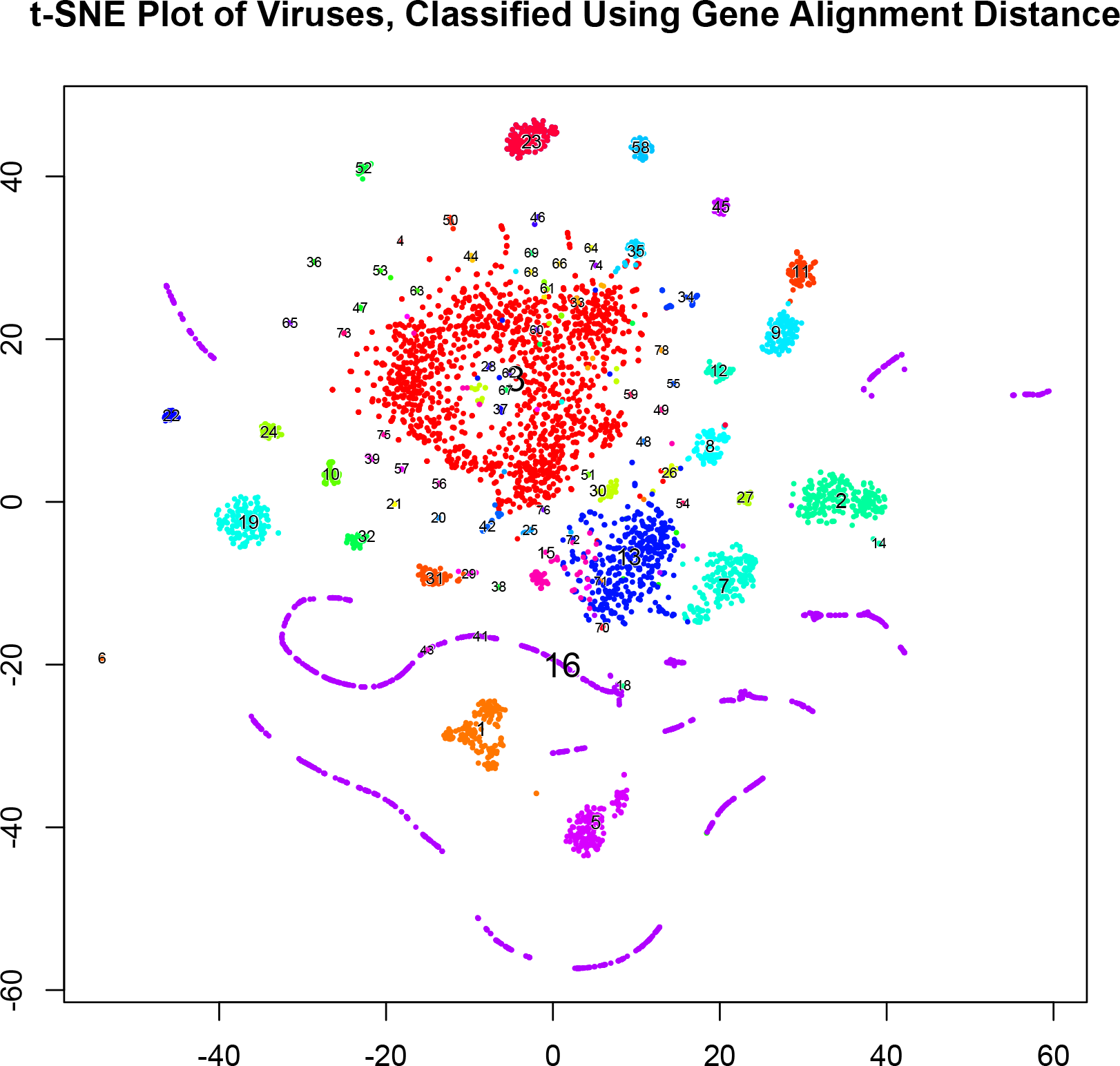
Supplementary plot of data classified using only the gene alignment distance. The cluster labels and colors are the same as in Figure 4. Notice that although many of the same structures are present, there is little distinction between smaller and larger groupings in the middle.

**Supplementary Figure 2.**
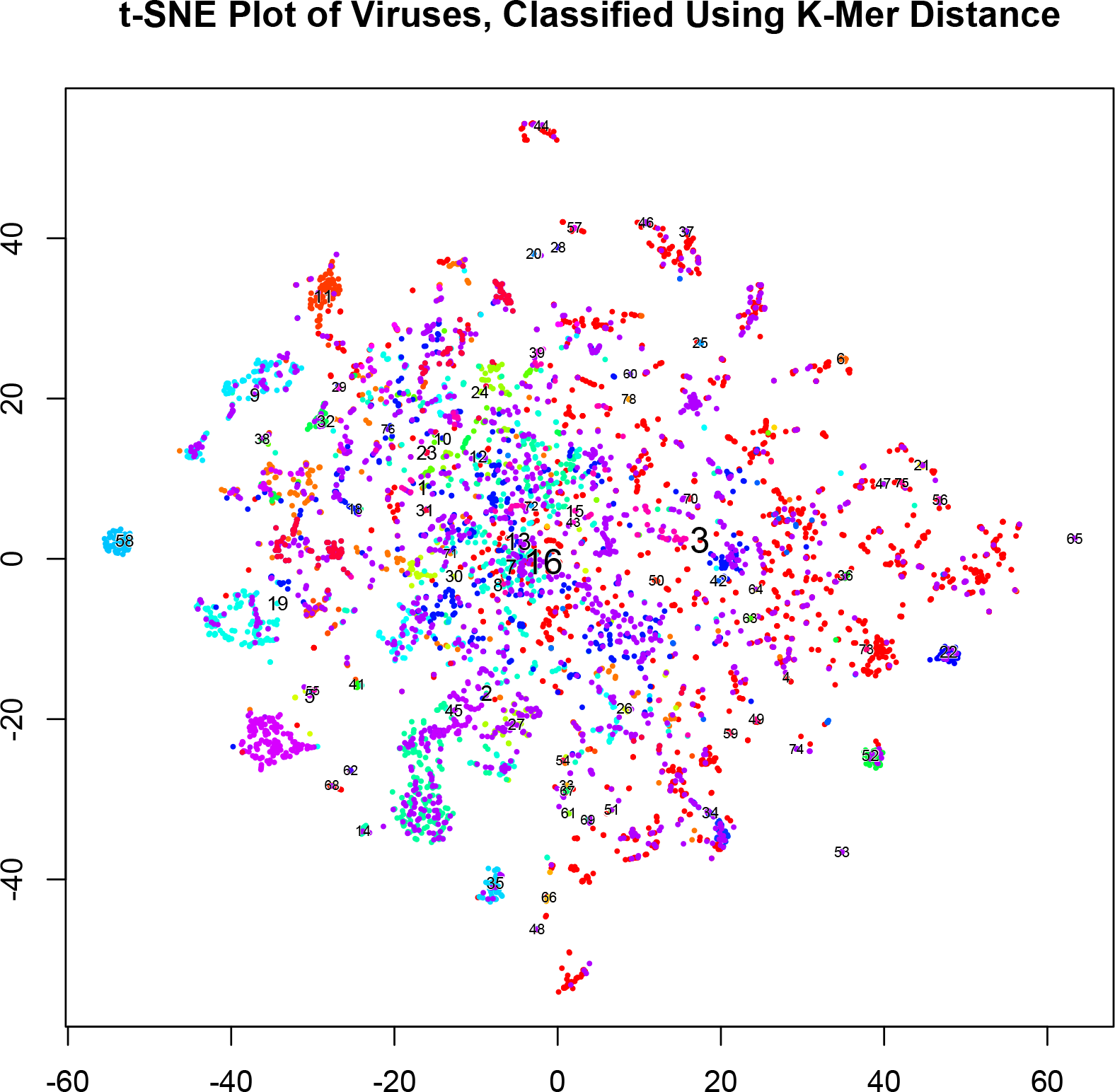
Supplementary plot of data classified using only the k-mer distance. The cluster labels and colors are the same as in Figure 4. Although the k-mer distance is able to provide some measure of similarity between very similar and very different sequences, it is not well-suited for an overall classification.

**Supplementary Figure 3.**
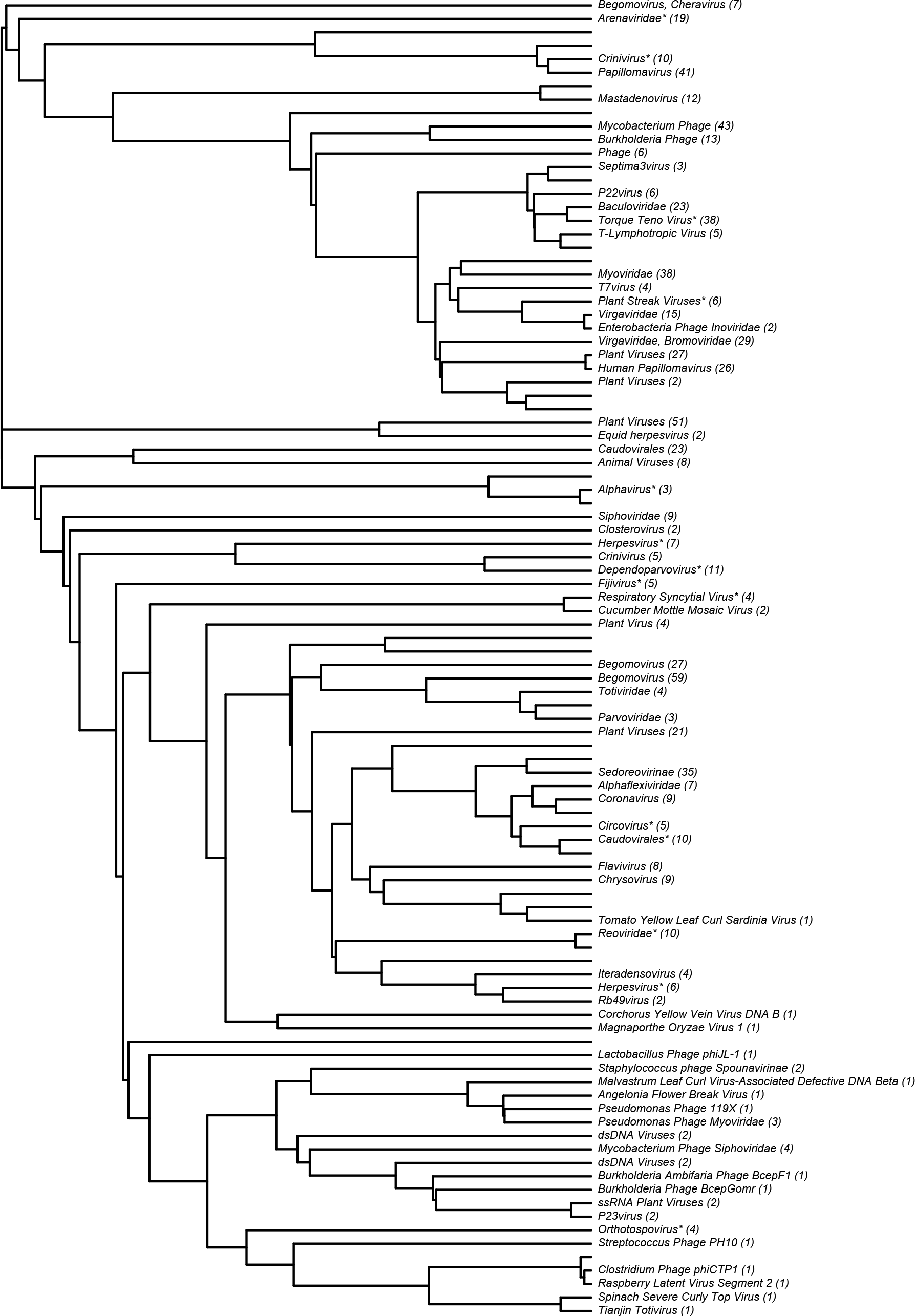
Dendrogram of 98 sub-clusters of cluster 16. The number of viruses in each cluster is given in parentheses. Relationships within sub-clusters are weaker, and a few clusters (those for which no name is given) displayed no unifying characteristics.

